# Ruminant-specific retrotransposons shape regulatory evolution of bovine immunity

**DOI:** 10.1101/2021.10.01.462810

**Authors:** Conor J. Kelly, Carol Chitko-McKown, Edward B. Chuong

## Abstract

Cattle are an important livestock species, and mapping the genomic architecture of agriculturally relevant traits such as disease susceptibility is a major challenge in the bovine research community. Lineage-specific transposable elements (TEs) are increasingly recognized to contribute to gene regulatory evolution and variation, but this possibility has been largely unexplored in ruminant genomes. We conducted epigenomic profiling of the type II interferon (IFN) response in bovine cells, and found thousands of ruminant-specific TEs including MER41_BT and Bov-A2 elements predicted to act as IFN-inducible enhancer elements. CRISPR knockout experiments in bovine cells established that critical immune factors including IFNAR2 and IL2RB are transcriptionally regulated by TE-derived enhancers. Finally, population genomic analysis of 38 individuals revealed that a subset of TE-derived enhancers represent polymorphic insertion sites in modern cattle. Our study reveals that lineage-specific TEs have shaped the evolution of ruminant IFN responses, and potentially continue to contribute to immune gene regulatory differences across modern breeds and individuals. Together with previous work in human cells, our findings demonstrate that lineage-specific TEs have been independently co-opted to regulate IFN-inducible gene expression in multiple species, supporting TE co-option as a recurrent mechanism driving the evolution of IFN-inducible transcriptional networks.

## INTRODUCTION

Modern cattle were domesticated 10,000 years ago and are an important livestock species (Troy et al. 2001; Orozco-terWengel et al. 2015). Cattle exhibit extensive phenotypic diversity, with over 1000 recognized breeds and closely related subspecies. Understanding the evolution and genetic basis of bovine biology, including traits involved in production and disease susceptibility, is a major challenge in the agricultural research community. However, most bovine genomic studies have focused on the effects of single nucleotide variants or small insertions and deletions (Littlejohn et al. 2016; Bouwman et al. 2018; Xiang et al. 2021). Transposable elements (TEs) could potentially have an important role shaping bovine variation and evolution, but with a few exceptions identified by genetic trait mapping (Liang et al. 2021; Trigo et al. 2021; Girardot et al. 2006; Albrecht et al. 2012; Schütz et al. 2016; Menzi et al. 2016), the impact of TEs on bovine biology remain largely unexplored.

Here, we investigated the significance of TEs on bovine biology, focusing on their capacity to act as gene regulatory elements. Epigenomic studies have revealed that TEs are a major source of cis-regulatory elements, and have been repeatedly co-opted to regulate the expression of host genes (Chuong et al. 2016; Fuentes et al. 2018; Ye et al. 2020). The cattle genome harbors numerous ruminant-specific retrotransposons including bovine endogenous retroviruses and the BovB retrotransposon (Garcia-Etxebarria and Jugo 2013; Ivancevic et al. 2018), which makes up over 25% of the cattle genome (Adelson et al. 2009). Ruminant-specific retrotransposons have recently been documented to show evidence of regulatory activity (Young et al. 2018; Halstead et al. 2020), but their consequences on bovine gene regulation are unknown.

To study the contribution of TEs to bovine gene regulation, we focused on the transcriptional network underlying the type II interferon response using interferon gamma (IFNG). We previously discovered that primate-specific TEs have been co-opted as enhancers that regulate antiviral IFNG-stimulated genes (ISGs) (Chuong et al. 2016). Given that innate immune responses show high interspecies variation (Shaw et al. 2017), we hypothesized that ruminant-specific TEs may have been independently co-opted to facilitate immune regulatory evolution in cattle. We conducted epigenomic profiling of the bovine IFNG response using MDBK cells, a model bovine cell line used to study innate immunity (Fay et al. 2020; Barreca and O’Hare 2004). We discovered that a substantial fraction of IFNG-inducible enhancers are derived from lineage-specific TEs, and CRISPR knockout studies revealed both bovine *IFNAR2* and *IL2RB* are regulated by TE-derived enhancers. Finally, we found that a subset of TE-derived enhancers exhibit insertional polymorphism in modern cattle. In the context of mammalian evolution, our study reveals that lineage-specific TEs have been repeatedly co-opted to regulate IFNG-stimulated genes in multiple species, and implicates TEs as an important source of regulatory variants affecting immune function.

## RESULTS

### Ruminant-specific retrotransposons contribute IFNG-inducible regulatory elements

To explore whether TEs contribute to bovine immune gene regulation, we conducted epigenomic profiling of the bovine type II IFN response (Fig 1A). We used RNA-seq to profile the transcriptional response to recombinant bovine IFNG in multiple cell types, including monocytes, leukocytes, B lymphocyte line BL3.1 cells, and MDBK cells. We identified a total of 4192 IFNG-stimulated genes (ISGs) showing inducible expression in at least one profiled cell type, with 298 shared across all cell types including canonical ISGs (Fig. 1B, 1C, Supplemental Fig. S1, Supplemental Table S1, S2). In parallel, we conducted chromatin profiling of IFNG-treated and untreated MDBK cells using ATAC-seq and CUT&RUN with antibodies against H3K27ac, phosphorylated RNA polymerase II subunit A (POL2RA), total STAT1, and phosphorylated STAT1 (Supplemental Table S3, S4). Successful antibody targeting of bovine STAT1 was validated by recovery of ISRE and GAS motifs in peaks called from IFNG-treated pulldowns (Supplemental Table S5).

**Figure 1.**
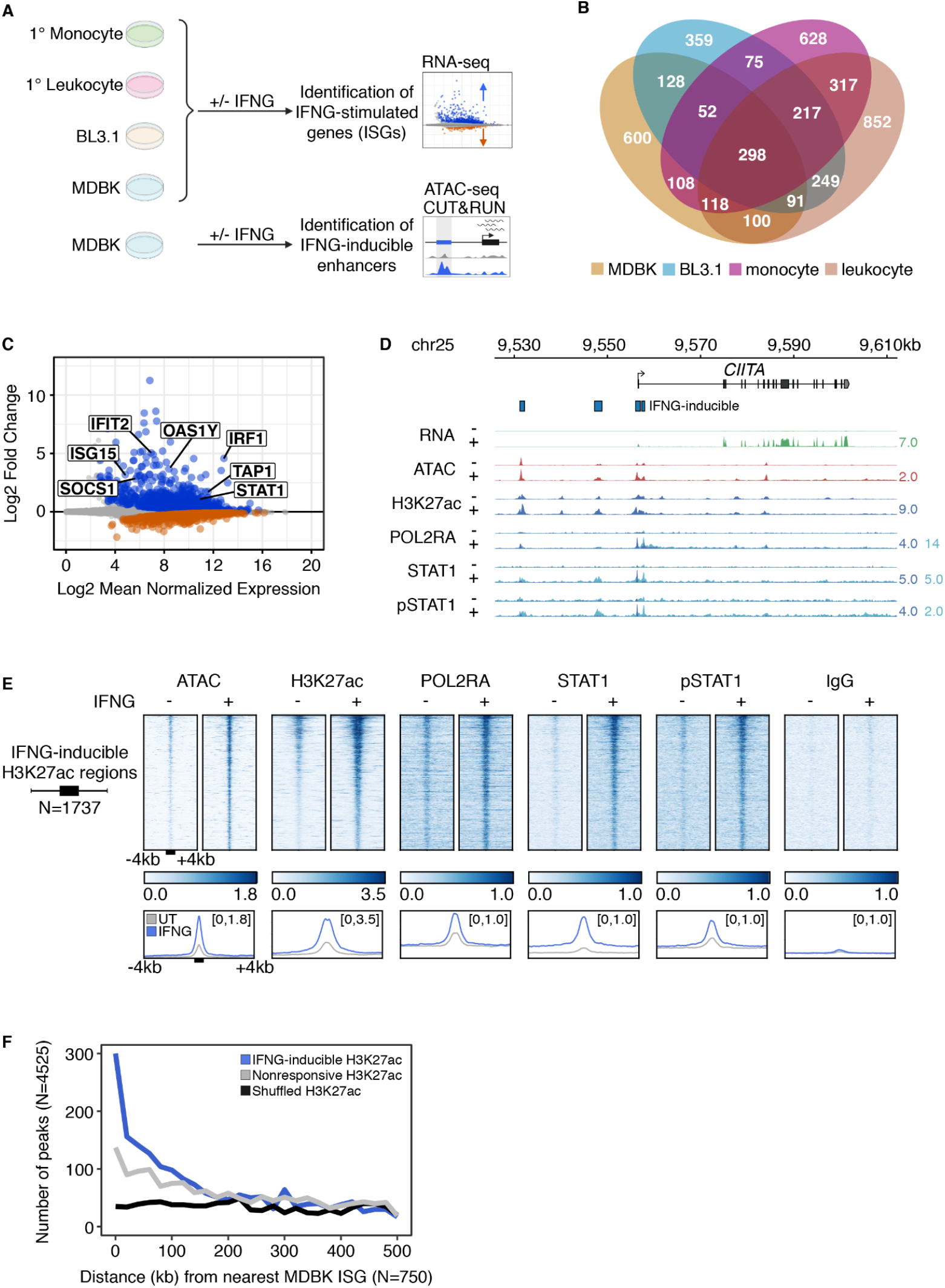
Epigenomic profiling of the bovine type II IFN response. (A) Schematic of experimental design. Made using Biorender. (B) Venn diagram of ISGs as defined by RNA-seq from MDBK, BL3.1, monocytes, and leukocytes. (C) MA plot of ISGs (blue) and IRGs (orange) from MDBK RNA-seq. Genes with an FDR < 0.05 are shown in grey, and canonical ISGs are labeled for reference. (D) Genome browser view of the *CIITA* locus. RNA-seq, ATAC-seq, and CUT&RUN tracks are normalized per million reads. CUT&RUN tracks for POL2RA, STAT1, and phosphorylated STAT1 pulldowns are divided by aligned fragments <= 150bp (dark blue) and > 150bp (light blue). The H3K27ac tracks correspond to all aligned fragments (dark blue). Predicted IFNG-inducible elements are shown. Signal track maxima are indicated on the right of each track. (E) Heatmaps showing normalized CUT&RUN signal (signal per million reads) over IFNG-inducible H3K27ac (N=1737) peaks. Bottom metaprofiles represent average normalized CUT&RUN signal across loci. (F) Frequency histogram of absolute distances from each peak set to the nearest ISG. ISG: Interferon-stimulated gene; IRG: Interferon-repressed gene.

We next used our CUT&RUN chromatin profiling data to predict the locations of IFNG-inducible enhancers, based on a significant increase in H3K27ac signal coverage. Examination of chromatin profiles at known ISGs such as *CIITA* and *TLR4* confirmed IFNG-inducible signal at predicted enhancers (Fig. 1D, Supplemental Fig. S2, S3). Using DESeq2, we identified 1737 elements with significantly increased H3K27ac read coverage across replicates (Supplemental Table S4). Consistent with IFNG-inducible enhancer activity, these elements displayed increased levels of chromatin accessibility, binding by total and phosphorylated STAT1, and POL2RA (Fig. 1E). These trends were not observed for elements with constitutive or decreased H3K27ac signal (Supplemental Fig. S4). Our set of predicted IFNG-inducible elements were also enriched for Gamma-interferon Activated Site (GAS) and IFN-Stimulated Response Element (ISRE) motifs predicted to bind STAT and IRF transcription factors (Supplemental Table S5), consistent with their activation by IFNG stimulation (Ivashkiv 2018). Inducible elements are also localized near ISGs compared to non-inducible enhancers or random genomic regions (Fig. 1F), showing strong enrichment within 20 kb of ISGs (*P* = 3.8⨯10^−300^) compared to non-responsive elements (*P* = 2.0⨯10^−58^) and randomly shuffled genomic regions (*P* = 0.72). These analyses indicate that our epigenomic dataset successfully captured inducible genes and regulatory elements underlying the type II IFN response in MDBK cells.

We next asked what fraction of predicted IFNG-inducible regulatory elements were derived from TEs. We defined IFNG-inducible regulatory elements based on H3K27ac signal, and used open chromatin peak summits to refine the location of regulatory elements due to the broad nature of H3K27ac enrichment (Methods). Out of 4198 IFNG-inducible regulatory elements, we found that 1658 (39.5%) have peak summits residing within ruminant-specific TEs (Fig. 2A, Supplemental Table S6), which is in line with previous epigenomic analyses in human and mouse cells (Chuong et al. 2016; Sundaram et al. 2014). In contrast, fewer (1065/5042, or 21.1%) of non-inducible or downregulated elements were derived from TEs, suggesting that TEs play a pronounced role in shaping the epigenome during immune stimulation. TE-derived inducible enhancers also showed strong colocalization with ISGs (Supplemental Fig. S6), consistent with a role in inducible gene regulation.

**Figure 2.**
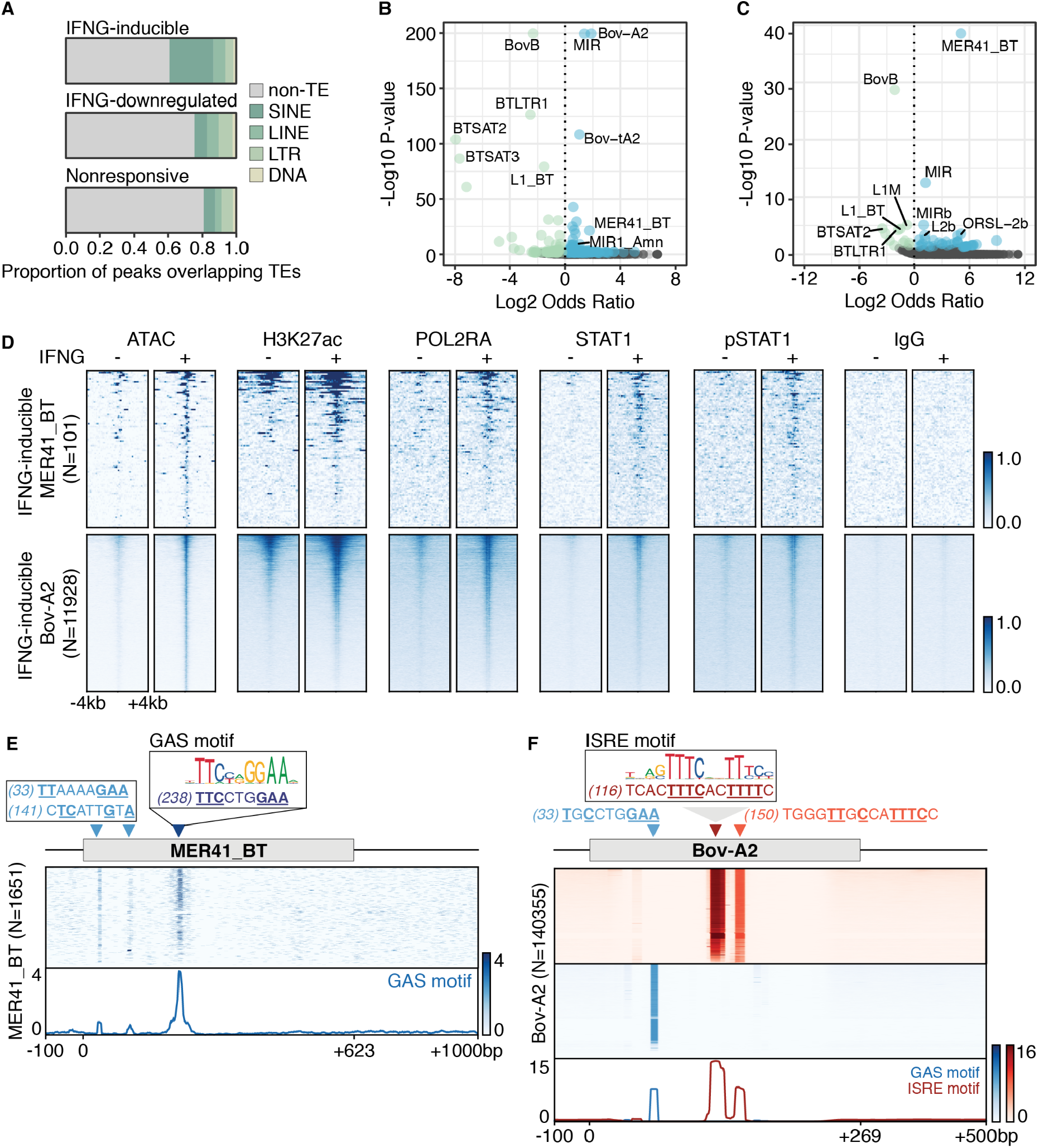
Contribution of ruminant-specific TEs to the IFNG-inducible regulatory landscape. (A) Proportion of IFNG-inducible (N=4198), IFNG-downregulated (N=1548), and nonresponsive H3K27ac peaks (N=3494) where the internal ATAC-seq summit overlaps a TE. (B) Volcano plot visualizing family-level TE enrichment for IFNG-inducible H3K27ac. TE families with a Fisher’s two-tail p-value < 0.05 were defined as enriched (blue) or depleted (green) based on the reported odds ratio. Nonsignificant families are shown in grey. (C) Family-level TE enrichment for IFNG-inducible STAT1 peaks, as in (B). (D) Heatmaps showing normalized ATAC-seq and CUT&RUN signal over IFNG-inducible MER41_BT (N=101, top) and Bov-A2 (N=11928, bottom) elements. (E) Schematic of IFNG-associated motifs within the MER41_BT consensus and extant sequences. Heatmap depicts the presence of motifs across 1651 extant MER41_BT elements, filtered for those that are at least 50% of full length relative to the consensus. Bottom metaprofile represents average signal across all elements. (F) Schematic of motifs as in (E) but across 140355 Bov-A2 elements, filtered for those within 5% of full length relative to the consensus. Heatmap intensity corresponds to motif matches based on the log likelihood ratio as defined by FIMO (Grant et al. 2011).

We next used Giggle (Layer et al. 2018) to determine whether any specific TE families were significantly overrepresented within the set of predicted IFNG-inducible enhancers (Methods). We found that the MER41_BT and Bov-A2 TE families were strongly overrepresented within regions showing IFNG-inducible levels of H3K27ac and STAT1 (Fig. 2B, 2C, Supplemental Fig. S7, Supplemental Table S7). At a genome-wide level, MER41_BT and Bov-A2 elements show inducible levels of H3K27ac, as well as increased binding of POL2RA and STAT1 (Fig 2D). These families contain perfect and partial matches to GAS or ISRE binding motifs within their consensus sequences and in most extant copies (Fig. 2E, 2F), indicating that MER41_BT and Bov-A2 elements are an abundant source of STAT1 binding sites in the bovine genome.

### Co-option of the MER41_BT elements to regulate IFNG-inducible immune gene expression

We next focused on investigating the regulatory impact of MER41_BT, a ruminant-specific ERV represented by 5491 elements in the cattle genome. 1399 of these contain at least one significant match to the GAS or ISRE motif (*P* < 1⨯10^−4^). Our epigenomic analysis of MDBK cells revealed that many MER41_BT elements showed canonical signs of IFNG-inducible enhancer activity, including inducible levels of H3K27ac, POL2RA, STAT1, and phosphorylated STAT1 (Fig 2D, Supplemental Fig. S6). A total of 101 MER41_BT elements show some evidence of inducible regulatory activity in MDBK cells, and 43 of which are located within 250 kb of an ISG (Supplemental Table S8). These observations suggest that a subset of MER41_BT elements may functionally contribute to the regulation of bovine ISGs.

We next asked whether individual MER41_BT elements function to regulate important immune genes in bovine cells. We first investigated an element located 41kb upstream of *IL2RB* (MER41_BT.IL2RB), which encodes a membrane-bound receptor for the interleukin-2 (IL2) and interleukin-15 (IL15) signaling pathways. *IL2RB* exhibits inducible expression in IFNG-stimulated MDBK cells and leukocytes, and LPS-stimulated bone marrow derived macrophages (BMDMs), suggesting that this element may regulate *IL2RB* (Fig. 3A, 3C). In addition to potentially acting as an enhancer for *IL2RB*, the MER41_BT element appears to act as a promoter of a bovine-specific transcript annotated as *LOC510185*. Consistent with the predicted inducible activity of the MER41 element, the *LOC510185* transcript exhibits IFNG-inducible expression in multiple bovine cell lines and primary cells, including monocytes, leukocytes, the BL3.1 B cell line, and MDBK cells. Additionally, *LOC510185* is expressed in BMDMs and alveolar macrophages infected with *M. bovis* (Fig. 3B) (Young et al. 2018; Hall et al. 2019). While the function of *LOC510185* is unknown, it arose from a recent tandem segmental duplication of the *IL2RB* locus, and its predicted open reading frame retains only the extracellular ligand binding domain of IL2RB (Supplemental Fig. S8). The widespread inducible expression of *LOC510185* in bovine cells suggests it may function as a bovine-specific factor involved in IL2 or IL15 signaling.

**Figure 3.**
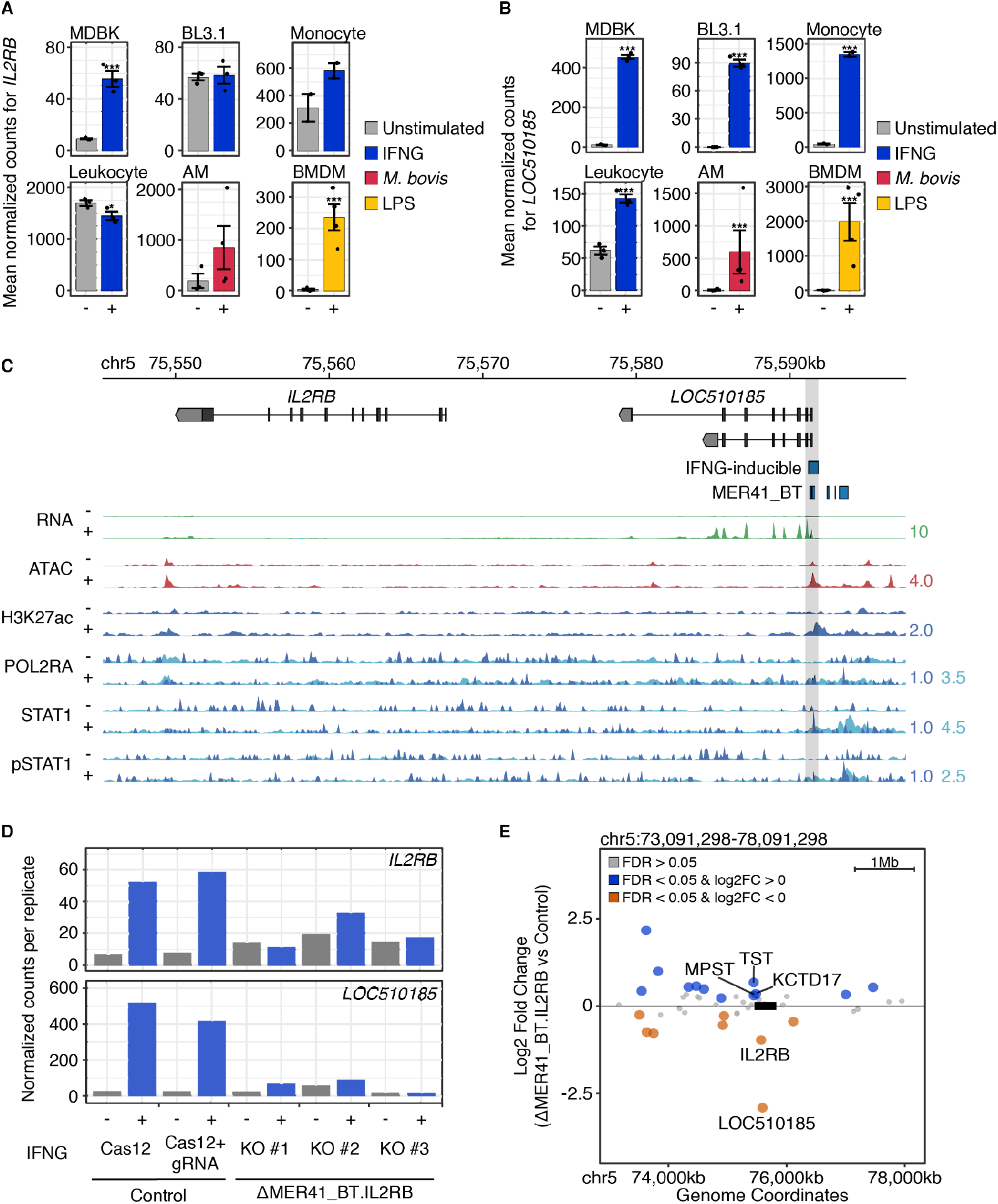
Co-option of a MER41_BT element for IL2RB and LOC510185 regulation. (A) DESeq2 normalized counts showing immune-stimulated expression for *IL2RB* from MDBK (N=3), BL3.1 (N=3), monocytes (N=2), leukocytes (N=3), alveolar macrophages (AM, N=3 untreated, N=4 stimulated), and bone marrow derived macrophages (BMDM, N=4). Treatments are indicated by color. ***: FDR < 0.0001, *: FDR < 0.01. Error bars denote SEM. (B) Normalized counts as in (A) but for *LOC510185*. (C) Genome browser view of *IL2RB* and *LOC510185*. RNA-seq, ATAC-seq, and CUT&RUN tracks are normalized per million reads. CUT&RUN signal profile tracks for POL2RA, STAT1, and pSTAT1 pulldowns were generated using sub-nucleosomal fragments <= 150bp (dark blue) as well as larger fragments > 150bp (light blue). The H3K27ac tracks correspond to all aligned fragments (dark blue). IFNG enhancers shown represent IFNG-inducible H3K27ac peaks that have been centered by ATAC-seq summits. MER41_BT.IL2RB is highlighted in grey. Values on the right of each track correspond to signal maxima. (D) Normalized DESeq2 counts for *IL2RB* (top) and *LOC510185* (bottom) from MDBK Cas12 controls and MER41_BT.IL2RB mutant clones. (E) Gene expression differences caused by the MER41_BT.IL2RB deletion within a 5 MB window centered on the deletion site (box not to scale), after IFNG-stimulation. Significantly upregulated (blue) and downregulated (orange) genes within 500 kb of the element are labeled. AM: alveolar macrophage; BMDM: bone marrow derived macrophage.

To experimentally characterize the potential promoter and enhancer activity of MER41_BT.IL2RB, we used CRISPR-Cas12 to delete the element in MDBK cells and determine its impact on gene expression. Although MDBK cells are an immortalized kidney cell line where *IL2RB* is only weakly expressed, transcriptomic evidence from primary immune cells suggests that inducible *IL2RB* and *LOC510185* expression is physiologically relevant. To specifically test regulatory activity, we generated deletions upstream of the predicted *LOC510185* transcriptional start site (Methods). We recovered three sequence-validated independent clones carrying homozygous deletions of MER41_BT.IL2RB (Supplemental Fig. S9), and profiled the IFNG responses for each clone using RNA-seq (Fig. 3D, Supplemental Table S9). In all clones, we observed a dramatic reduction in IFNG-stimulated expression levels of both *IL2RB* (FDR = 1.1×10^−3^) and *LOC510185* (FDR = 1.7×10^−10^) (Fig 3E). Together, these results show that a specific MER41_BT element functions simultaneously as a IFNG-inducible promoter of *LOC510185* and an IFNG-inducible enhancer of *IL2RB* in MDBK cells.

We next characterized a separate MER41_BT element located 19 kb upstream of the gene *IFNAR2* (MER41_BT.IFNAR2), which encodes a membrane-bound receptor mediating Type I interferon signaling. Bovine *IFNAR2* shows IFNG-inducible expression in monocytes and leukocytes, MDBK cells, and BL3.1 cells (Fig. 4A). Additionally, *IFNAR2* expression is induced in alveolar macrophages challenged with *M. bovis* and LPS-stimulated BMDMs. The predicted upstream MER41_BT.IFNAR2 element shows inducible enhancer activity in MDBK cells (Fig. 4B). To test whether this element regulates bovine *IFNAR2*, we used CRISPR-Cas12 to delete a region containing GAS binding motifs within the MER41_BT.IFNAR2 element (Supplemental Fig. S10). We isolated 2 independent clones carrying the expected deletion and profiled the IFNG response in each clone using RNA-seq (Fig 4C, Supplemental Table S9). We found that *IFNAR2* was still inducible by IFNG stimulation, but showed significantly reduced expression compared to control cells (FDR = 2.4⨯10^−6^) (Fig. 4C, 4D). We also found that the nearby interferon gamma receptor gene *IFNGR2* showed significantly reduced expression under IFNG stimulation (FDR = 3.0⨯10^−6^) (Fig. 4D, Supplemental Fig. S11), indicating that deletion of the MER41_BT element affected the expression of multiple genes. We did not observe complete ablation of inducible *IFNAR2* or *IFNGR2* expression, potentially due to the presence of nearby IFNG-inducible enhancers that could compensate for the deletion (Fig 4B). These results demonstrate that MER41_BT.IFNAR2 functions as an enhancer that contributes to IFNG-inducible regulation of *IFNAR2* and *IFNGR2* in MDBK cells.

**Figure 4.**
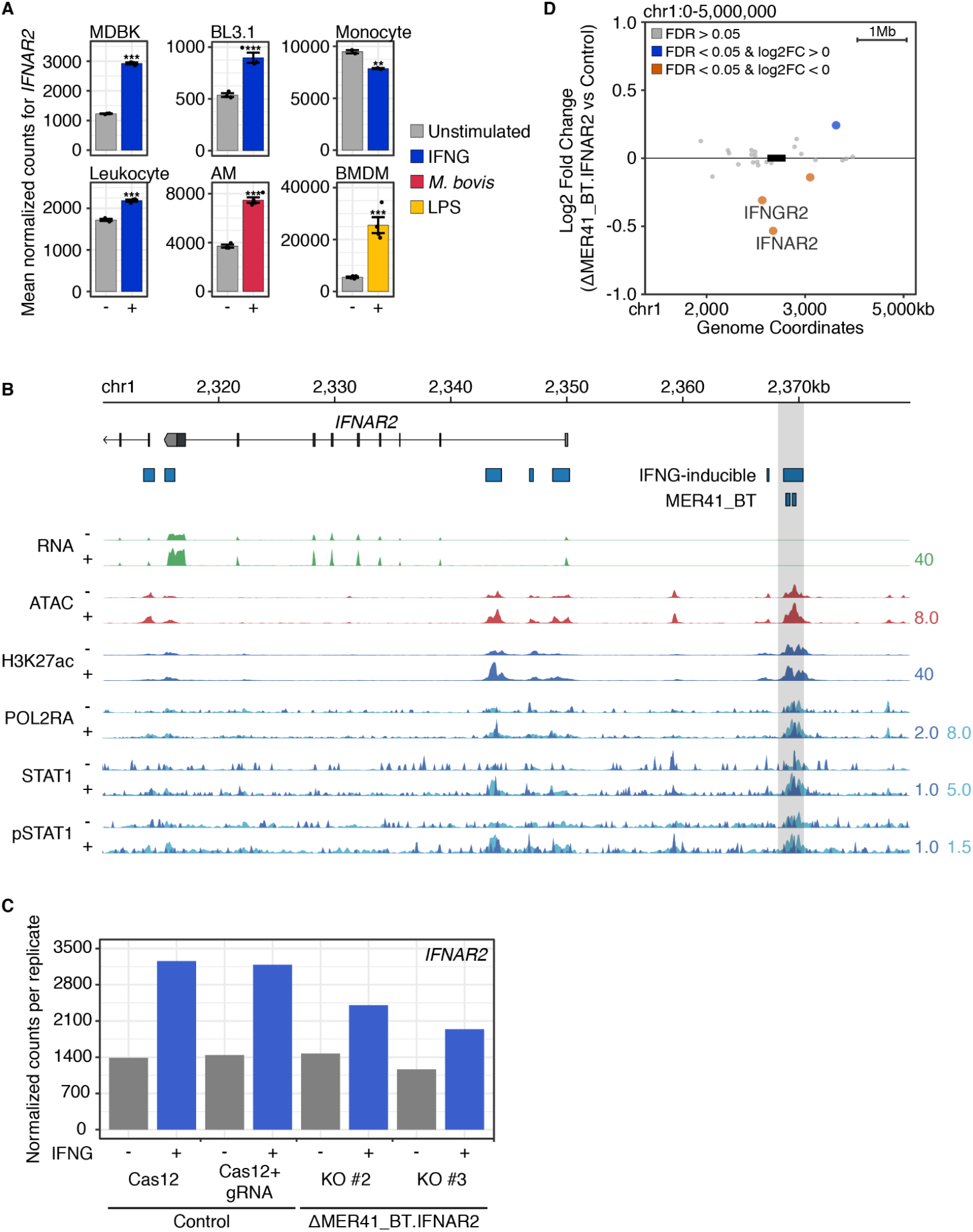
Co-option of a MER41_BT element for IFNAR2 regulation. (A) Mean DESeq2 normalized counts showing expression for *IFNAR2* from wild type bovine cells. ***: FDR < 0.0001, **: FDR < 0.001. Error bars denote SEM. (B) Genome browser view of *IFNAR2*. RNA-seq, ATAC-seq, and CUT&RUN tracks are normalized per million reads. CUT&RUN tracks for POL2RA, STAT1, and pSTAT1 pulldowns are divided by aligned fragments <= 150bp (dark blue) and > 150bp (light blue). The H3K27ac tracks correspond to all aligned fragments (dark blue). IFNG enhancers shown indicate predicted IFNG-inducible H3K27ac peaks. MER41_BT.IFNAR2 is highlighted in grey. Values on the right of each track correspond to signal maxima. (C) Normalized DESeq2 counts for *IFNAR2* from MDBK Cas12 controls and MER41_IFNAR2 mutant clones. (D) Gene expression differences caused by the MER41_BT.IFNAR2 deletion within a 5 MB window centered on the deletion site (box not to scale), after IFNG stimulation. Significantly upregulated (blue) and downregulated (orange) genes within 500 kb of the element are labeled. AM: alveolar macrophage; BMDM: bone marrow derived macrophage.

### TEs contribute to evolutionary diversification of ruminant immune responses

Having established that a subset of ruminant-specific TEs function to regulate ISGs in bovine cells, we next examined their impact on ruminant genome evolution. The MER41_BT family was originally annotated as a bovine-specific ERV belonging to the MER41-like family of ERVs, which are present in several other lineages including primates and carnivores (Bao et al. 2015). MER41-like ERVs from different lineages exhibit high sequence similarity, consistent with independent germline integrations by a retroviral lineage that underwent multiple cross-species transmissions (Zhuo and Feschotte 2015). To better resolve the timing of MER41_BT emergence, we re-annotated repeats in the assembled reference genomes of 30 mammals including 19 cetartiodactyl species (Supplemental Table S10), using a uniform set of cetartiodactyl MER41-like consensus sequences (Methods). Based on the presence and abundance in each species clade, we estimated when each cetartiodactyl MER41 subfamily originated (Fig. 5A). This confirmed that primate MER41 and bovine MER41_BT arose from separate germline integration events, and revealed that MER41_BT elements are present in pigs and all cetruminantia species. MER41_BT elements derive from an ancestral MER41-like retrovirus that integrated into an ancestral cetartiodactyl genome roughly 70 mya, prior to the divergence of camelidae and artiofabula (Fig. 5B). This estimate coincides with the emergence of primate MER41 elements after the divergence of anthropoid primates from prosimian lineage roughly 70 mya, consistent with a scenario where primate and ruminant MER41 derive from separate integrations by a common ancestral retroviral lineage.

**Figure 5.**
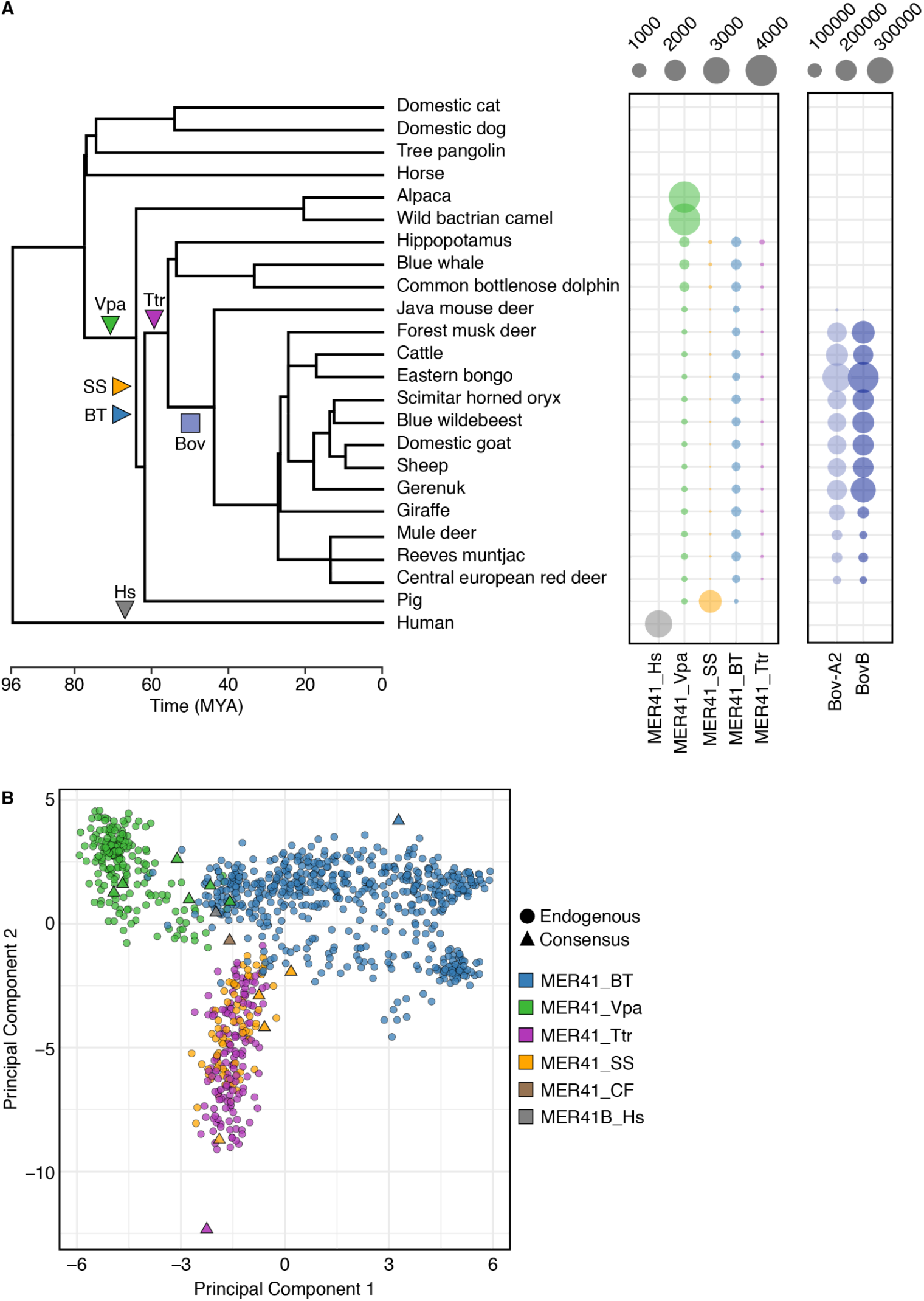
Mapping TE subfamily evolution in cetartiodactyla. (A) Number of MER41, Bov-A2, and BovB elements identified in 24 mammalian assemblies using RepeatMasker. Bubble size corresponds to the number of TEs annotated in each assembly. Phylogeny was obtained from TimeTree (Kumar et al. 2017) and depicts species divergence. Estimated insertion times are indicated on the phylogeny as triangles (MER41) and a square (Bov-A2 and BovB). (B) PCA plot of endogenous bovine MER41 elements (circles) and cetartiodactyl, carnivore (MER41_CF), and human (MER41B_Hs) MER41 consensus sequences (triangles) colored by subfamily. All elements were aligned using MUSCLE (Edgar 2004), and transformed values indicating similarity were obtained using Jalview (Waterhouse et al. 2009). Consensus sequences are named based on the species in which they were discovered. Hs: *Homo sapiens*; Vpa: *Vicugna pacos*; Ttr: *Tursiops truncatas*; SS: *Sus scrofa*; CF: *Canis familiaris*; BT: *Bos taurus*.

We next used synteny analysis to determine whether individual MER41_BT insertions were conserved across cetruminantia species. We asked whether each annotated element in the cattle genome, along with flanking sequence, was present in the genomes of select cetartiodactyl species. This revealed that the majority of insertions were poorly conserved across cetruminant species (Supplemental Table S11), suggesting that MER41_BT continued to propagate in host genomes as cetruminant lineages diverged. Given our evidence that MER41_BT influences gene regulation in cattle, it is possible that lineage-specific insertions of MER41_BT elements have contributed to gene regulatory diversification across cetruminant lineages.

Next, we examined the evolutionary history of Bov-A2 elements, which our analysis identified as another major source of predicted IFNG-inducible enhancers. Bov-A2 elements are SINE-like elements derived from the BovB retrotransposon roughly 20-30 mya and constitute one of the most abundant sequences in bovine genomes (Nijman et al. 2002; Nilsson et al. 2012). There are over 360,000 Bov-A2 sequences in the cattle genome, 280,000 of which contain at least one GAS or ISRE motif. In MDBK cells, Bov-A2 and other BovB derivatives represent 24.3% of all predicted IFNG-inducible regulatory elements, or 61.6% of all TE-derived inducible elements (Supplemental Table S6). Compared to MER41_BT, Bov-A2 elements were active more recently (Nijman et al. 2002; Nilsson et al. 2012; Dekel et al. 2015; Young et al. 2018), but no study to date has directly examined whether Bov-A2 insertions are polymorphic within modern species.

We investigated whether Bov-A2 elements are a source of modern variation by using MELT (Gardner et al. 2017) to identify polymorphic mobile element insertions in a diverse panel of 38 cattle whole-genome sequences (Workman et al. 2018; Tsuda et al. 2013; Stothard et al. 2015; Shin et al. 2014; Bickhart et al. 2016; Heaton et al. 2016; Chen et al. 2018). Across all individuals, our analysis revealed 1017 Bov-A2 elements with read-supported evidence for insertional polymorphism, including 740 non-reference insertions and 277 deletions relative to the reference Hereford *bosTau9/ARS-UCD1*.*2* assembly (Fig. 6A, Supplemental Fig. S12A, Supplemental Table S12). We also observed 1782 polymorphic BovB elements, which are the proposed originator of Bov-A2 elements (Fig. 6A, Supplemental Table S12) (Nijman et al. 2002; Nilsson et al. 2012). As expected, we did not identify any insertional polymorphisms of the more ancient MER41_BT family (Fig. 6A, Supplemental Table S12). Many Bov-A2 and BovB polymorphisms were present in more than one individual (Fig. 6B) and principal components analysis of cattle genomes based on Bov-A2 and BovB insertion status corresponded with subspecies (Fig. 6C, Supplemental Fig. 12B). Collectively, our analysis indicates that Bov-A2 insertions are a current source of structural genetic variants in modern cattle individuals.

**Figure 6.**
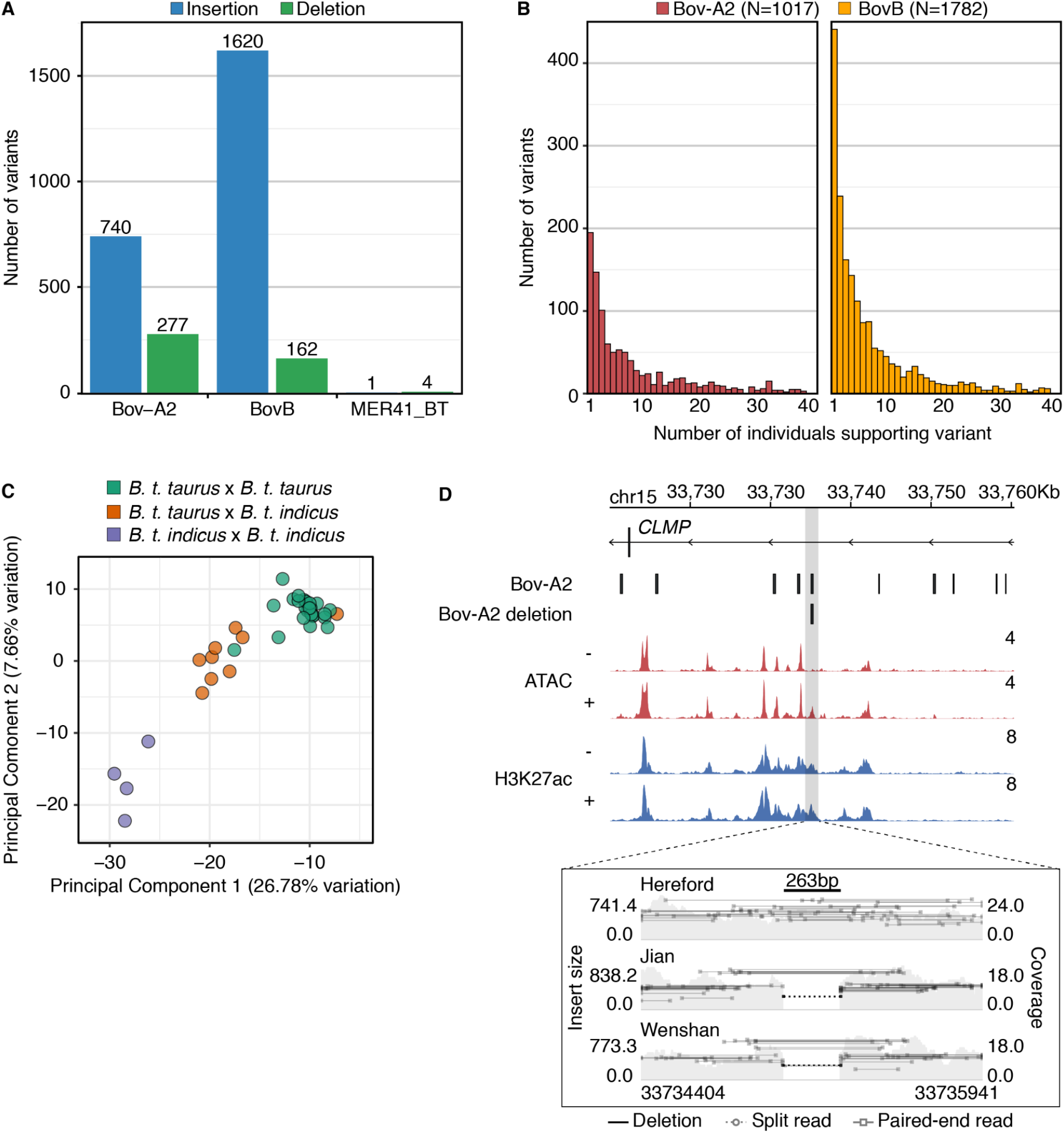
Detection of Bov-A2 insertion polymorphisms. (A) Number of polymorphic Bov-A2, BovB, and MER41_BT insertions (blue) and deletions (green) for each TE family as called by MELT (Gardner et al. 2017). (B) Histogram of polymorphic Bov-A2 variants as a function of number of individuals supporting each variant. Individuals were defined as supporting a variant if they showed evidence of at least one variant allele. (C) Histogram of polymorphic BovB variants as a function of number of individuals supporting each variant. Individuals were called as supporting a variant if they showed evidence of at least one variant allele. (D) Genome browser view (top) of a putative Bov-A2 deletion. ATAC-seq and CUT&RUN tracks from MDBK are normalized per million reads. Values on the right of each track correspond to signal maxima. Variant visualization plot (bottom) was produced using Samplot (Belyeu et al. 2021) and depicts aligned fragments from three individuals over the predicted Bov-A2 deletion.

Finally, we asked whether any polymorphic Bov-A2 insertions show evidence of regulatory activity. Based on our epigenomic data from MDBK cells, we identified 17 Bov-A2 elements with regulatory activity that were present in the Hereford reference genome and MDBK cells, but missing in at least one allele in our analysis (Supplemental Table S13). This included a Bov-A2-derived element located near the bovine gene *CLMP*, which was present in the Hereford reference genome and MDBK cells, but missing in Angus individuals (Fig. 6D). Additionally we identified polymorphic elements in the first intron of *BICC1* and 26 kb upstream of *SNX19* (Supplemental Fig. 12C, 12D). While more work is necessary to causally link Bov-A2 polymorphic loci to functional variation, our observations suggest that TE insertional polymorphisms may be an underappreciated class of regulatory variants that contribute to differences in immune gene expression.

## DISCUSSION

Infectious diseases have a large economic impact on the cattle industry, and understanding the genetic architecture of bovine immune responses is key for improving treatments and breeding by genomic selection (Mallard et al. 2015; Keirn et al. 2001; Stear et al. 2001; Thompson-Crispi et al. 2014). To this end, thousands of individual genomes from diverse cattle breeds and other ruminant species have been sequenced to date (Hayes and Daetwyler 2019). However, the majority of association studies have focused on SNPs, with structural variants only recently gaining attention (Liu et al. 2010; Hu et al. 2020; Chen et al. 2017). Lineage-specific TEs are a prominent source of genomic variation that have been implicated in trait diversity in many domesticated species (Langevin et al. 2018; Bannasch et al. 2021; Lisch 2013; Wragg et al. 2013), but there are currently only a handful of examples of TE insertions affecting ruminant variation, including cholesterol deficiency and coat color (Menzi et al. 2016; Liang et al. 2021). Our epigenomic approach uncovered TEs as a source of IFNG-inducible enhancers that regulate bovine ISGs, with some that are still polymorphic in modern cattle. Our findings warrant further examination of TE insertions as a class of variants to be considered in bovine genome-wide association studies, potentially by adopting new graph-based reference genomes that encompass bovine structural variants (Crysnanto et al. 2021).

By investigating TE-mediated regulation in a less well-studied organism, our study also uncovered a striking example of parallel evolution, involving the independent co-option of the MER41 family of retroviruses in both primates and ruminants. Primate MER41 and ruminant MER41_BT share high sequence identity, but originate from independent germline integration events, consistent with separate germline integrations by a retroviral lineage that underwent multiple cross-species transmissions (Zhuo and Feschotte 2015). Therefore, we speculate that descendents of an ancestral MER41 retrovirus lineage integrated into the genomes of multiple mammalian lineages in a time period of 60-70 mya, including primates and cetartiodactyla. Remarkably, our study found that MER41 elements integrated into separate lineages have been convergently co-opted for similar regulatory functions regulating host IFNG responses. It is possible that the sequences of ancient MER41 retroviruses may have had regulatory properties, such as inducible activation, that predisposed them to successful endogenization and eventual domestication for host function.

Our study also revealed pervasive enhancer activity at Bov-A2 elements, which are amongst the most abundant sequences in cattle genomes. Bov-A2 elements are SINEs derived from the BovB retrotransposon, which integrated into ruminant genomes via horizontal transfer from a reptile (Ivancevic et al. 2018). Individual Bov-A2 elements have previously been characterized as a source of LPS-inducible promoter sequences (Young et al. 2018), and our study reveals that thousands of Bov-A2 elements may possess IFNG-inducible regulatory activity. Together with our analysis confirming that Bov-A2 insertions are still polymorphic in modern cattle, these findings implicate an important role for TEs in contributing to both ruminant evolution and modern genetic variation. While further experimental investigation will be necessary to link TE-derived regulatory variants to phenotypic differences, our findings suggest that TEs may underlie differences in immune gene regulation across bovine individuals and populations.

Together with our previous work in human cells (Chuong et al. 2016), our discovery that lineage-specific TEs have been independently co-opted to regulate ISGs in multiple species draws parallels to the finding that ERVs have been repeatedly co-opted to mediate placentation in mammals (Dupressoir et al. 2012; Sun et al. 2021; Chuong et al. 2013; Dunn-Fletcher et al. 2018). A growing number of case examples have been discovered where unrelated TEs have been co-opted for convergent functions in multiple species, suggesting that TEs may have a propensity to facilitate the evolution of certain biological processes, particularly those involving genetic conflicts. Studies investigating the epigenetic regulation of IFN responses diverse vertebrate species may therefore uncover a widespread role for TEs in mediating the evolution of innate immune regulatory networks.

## METHODS

### Reagents and sequences

For a detailed list of all reagents, primer sequences, and gRNA sequences used in this study see Supplementary Table S14.

### Cell lines

Bovine MDBK cells (gift from Sara Sawyer) were cultured in minimum essential medium (MEM) supplemented with nonessential amino acids, 1X penicillin-streptomycin, and 10% fetal bovine serum. MDBK cells were routinely passaged using accutase. BL3.1 cells were cultured in Roswell Park Memorial Institute (RPMI) 1640 medium supplemented with 1X penicillin-streptomycin and 10% fetal bovine serum. All IFNG treatments were performed using 100 ng/mL recombinant bovine IFNG. Both cell lines were cultured at 37°C and 5% CO_2_.

### Isolation of primary monocyte and leukocytes

Monocytes and leukocytes were derived from peripheral blood mononuclear cells (PBMCs) from different individuals. 15 mL of blood were obtained from two male cattle by jugular venipuncture into 60 cc heparinized syringes. Both sampled animals are referred to as “line 999” and exhibit the following breed makeup: 1/2 Angus, 1/4 Gelbvieh, 3/16 Charolais, 1/16 Hereford. Monocytes and leukocytes were extracted from a 8.5 month old cryptorchid bull and 11 month old steer, respectively. Animal handling and isolation of bovine blood cells was approved by the U.S. Meat Animal Research Center (USMARC) Institutional Animal Care and Use Committee (#2.1).

15 mL of blood was combined with 15 mL 1X PBS and 15 mL Ficoll-Paque Plus and centrifuged for 45 minutes at 1,000 x g at room temperature. The PBMC layer was retained, washed in 50 mL of cold 1X PBS, and lysed in 1 mL Red Blood Cell Lysis Buffer for 5 minutes at 37°C and 5% CO_2_. The PBMCs were washed three times with 50 mL, 10 mL, and 10 mL cold 1X PBS, respectively, and resuspended in 45 mL of FBS-free RPMI 1640. Monocytes were isolated as described in (Chitko-McKown et al. 2004). PBMCs were added to treated T75 flasks and incubated at 37°C and 5% CO_2_ overnight, and the non-adherent, predominantly leukocyte fraction was removed by washing with PBS. The remaining adherent population represents the monocyte fraction. Leukocytes were derived from the non-adherent fraction. IFNG treatments were performed with the PBMC fraction prior to separating adherent and non-adherent populations.

### Lentivirus production

24 h prior to transfection 4⨯10^6^ viable HEK293T cells were seeded in a coated 10cm dish in 10 mL Dulbecco’s modified Eagle’s medium (DMEM), 1X penicillin-streptomycin, and 10% fetal bovine serum. Prior to transfection the media was replaced with antibiotic-free DMEM supplemented with 10% fetal bovine serum. For transfection FuGENE was used as follows: 20 ug psPax2, 2 ug pCMV-VSV-G, 16 ug pRDA_174 or pRDA_052_crRNA, 114 uL FuGENE, and Opti-MEM (up to 2.5 mL) were combined and incubated at room temperature for 20 minutes. The mixture was added dropwise to the dish containing HEK293T cells, and the cells were incubated at 37°C and 5% CO_2_. After 6 h the media was replaced with DMEM supplemented with 1X penicillin-streptomycin and 10% fetal bovine serum. Virus was harvested 36 h after this media change and filtered using a 0.45 um PES filter.

### Generation and analysis of CRISPR-enAsCas12a mutants

To generate an MDBK line that stably expresses enAsCas12a, the following procedure was used: 24 h prior to transduction, 1⨯10^5^ viable MDBK cells were seeded into coated 6-well plates. Before transduction the media in each well was swapped with MEM supplemented with 10% fetal bovine serum and 8 ug/mL polybrene for a final volume of 2 mL after the addition of lentivirus. Each well was transduced with different volumes of pRDA_174 lentivirus (50, 200, 600, 1000 uL). After 24 h the media was replaced with fresh complete MEM. 24 h after this media change 4 ug/mL blasticidin was added to each well to select for cells stably expressing enAsCas12a. This selection continued for a total of 14 days, after which one well (200 uL) was chosen for further use. The presence of integrated enAsCas12a was validated by gDNA PCR. Clonal lines were isolated using array dilution method with 25% conditioned MEM, and 6 clones were screened for expression of integrated enAsCas12a by RT-qPCR. One clone was selected for further experimentation and maintained under 2 ug/mL blasticidin selection.

To generate each MER41 deletion, MDBK cells stably expressing enAsCas12a were transduced with pRDA_052_cRNA lentivirus containing target gRNAs, following the same procedure above. 48 h after transduction each well was treated with 2 ug/mL puromycin. Puromycin selection continued for a total of 7 days. Evidence of editing was confirmed using gDNA PCR. Cells that were transduced with 200 uL lentivirus were clonally expanded using the array dilution method with 25% condition MEM, and 80-120 clones were screened for homozygous deletions by PCR using both internal and flanking primer pairs at the expected deletion site. To determine deletion breakpoint sequences, PCR products flanking each deletion site were cloned into pJET and transformed into NEB 5-alpha Competent *E. coli*, and at least 2 individual colonies were sequenced using Sanger sequencing.

### Preparation of gRNA constructs

For each MER41 element (associated with *IFNAR2* and *LOC510185*), two 23bp gRNA sequences were designed to generate a single internal deletion that encompasses the putative STAT1 binding sites. Each pair of gRNAs was synthesized as a single crRNA oligonucleotide and included BsmBI sites with accompanying overlaps (bolded sequences) as explained in (DeWeirdt et al. 2021): 5’-[handle]CGTCTCA**AGAT**[gRNA1][DR-wt][gRNA2][DR-1]TTTTTT***GAAT***CGAGACG[handle]

Sequence was added to the 5’ and 3’ ends to help facilitate BsmBI recognition and to meet the 300bp requirement for synthesis through Twist Bioscience. Only gRNA sequences with a first percentile (see (DeWeirdt et al. 2021) 5’ PAM were considered. All gRNA sequences were verified to uniquely target the locus of interest using BLAT (Kent 2002) against the bosTau9 assembly. crRNA oligonucleotides were synthesized by Twist Bioscience as a single, 300bp oligonucleotide or precloned into pTwist-Amp-High.

Each crRNA oligonucleotide was cloned into pRDA_052 following a modified protocol from (DeWeirdt et al. 2021). Briefly, crRNA oligonucleotides were cloned into pRDA_052 using 0.10 fmol crRNA oligonucleotide, 0.05 fmol pRDA_052, 2.00 uL T4 DNA Ligase Buffer, 1.00 uL T4 DNA Ligase, 1.00 uL BsmBI-v2, and nuclease-free water up to 20 uL. crRNA olignonucleotides that were precloned into pTwist-Amp-High were subcloned into pRDA_052 using 75 ng pTwist-Amp-High-crRNA, 75 ng pRDA_052, 2.00 uL T4 DNA Ligase Buffer, 1.00 uL T4 DNA Ligase, 1.00 uL BsmBI-v2, and nuclease-free water up to 20 uL. PCR cycling conditions: 60 minutes at 42°C, 5 minutes at 60°C, and 10 minutes at 65°C. 2 uL of each ligation reaction was used to transform NEB Stable Competent *E. coli*. Plasmid DNA was harvested using Zymo ZymoPURE II Plasmid Midiprep Kit, and the sequence of each construct was verified by Sanger sequencing (Quintara Biosciences, Fort Collins, CO and Genewiz, South Plainfield, NJ).

### ATAC-seq

ATAC-seq libraries were prepared following (Corces et al. 2017) using 5×10^4^ viable (>90%) MDBK cells. 30 minutes before harvesting, cells were treated with 200 U/mL DNase I for 30 minutes. Each 5⨯10^4^ sample was transposed using 100 nM transposase as specified in (Corces et al. 2017). Preamplifcation was performed by combining 20 uL transposed sample, 2.5 uL 25 uM i5 primer, 2.5 uL 25 uM i7 primer, and 25 uL 2x NEBNext Ultra II Q5 Master Mix. PCR cycling conditions: 30 s at 98°C, 5x(10 s at 98°C, 30 s at 63°C, 45 s at 65°C). qPCR was performed as follows: 5 uL of preamplified sample was combined with 0.5 uL 25 uM i5 primer, 0.5 uL 25 uM i7 primer, 3.75 uL nuclease-free water, 0.25 uL 25X SYBR Gold, and 5 uL 2x NEBNext Ultra II Q5 Master Mix. qPCR cycling conditions: 30 s at 98°C, 20x(10 s at 98°C, 30 s at 63°C, 45 s at 65°C). Preamplified samples were amplified for another 7 cycles. Final libraries were cleaned using a 0.5X/1.3X double-sided bead cleanup using KAPA Pure Beads and quantified using Qubit dsDNA High Sensitivity and TapeStation 4200 HSD5000. Libraries were pooled and sequenced on an Illumina NovaSeq 6000 (University of Colorado Genomics Core) as 150bp paired-end reads.

### CUT&RUN

CUT&RUN pulldowns were generated using a protocol from (Janssens and Henikoff 2019; Meers et al. 2019). All buffers were prepared according to the “High Ca^2+^/Low Salt” section of the protocol using 0.04% digitonin. 5⨯10^5^ viable cells were used for each pulldown. The following antibodies were used: rabbit anti-mouse IgG (1:100), rabbit anti-H3K27ac (1:100), rabbit anti-pRPB1-Ser5 (1:50), rabbit anti-STAT1 (1:100), rabbit anti-pSTAT1-Ser727 (1:100). pA-MNase (gift from Steve Henikoff) was added to each sample following primary antibody incubation at a final concentration of 700 ng/mL. Chromatin digestion, release, and extraction was carried out according to (Janssens and Henikoff 2019; Meers et al. 2019). Pulldown success was determined by Qubit dsDNA High Sensitivity and TapeStation 4200 HSD5000 before proceeding with library preparation.

Libraries were generated using a modified protocol for use with the KAPA HyperPrep Kit. Briefly, the full volume of each pulldown (∼30 uL) was diluted to 50 uL in nuclease-free water, and libraries were generated following the manufacturer’s protocol with the following modifications. Freshly diluted 0.200 uM single-index adapters were added to each library at a low concentration (9 nM) to minimize adapter dimer formation. Adapter-ligated libraries were treated with 0.2% SDS and 0.4 mg/mL Proteinase K for 1 h at 37°C and underwent a double-sided 0.8X/1.0X cleanup with KAPA Pure Beads. Purified, adapter-ligated libraries were amplified using the following PCR cycling conditions: 45 s at 98°C, 15x(15 s at 98°C, 10 s at 60°C), 60 s at 72°C. Amplified libraries underwent a double-sided 0.8X/1.0X cleanup. The final libraries were quantified using Qubit dsDNA High Sensitivity and TapeStation 4200 HSD5000. Libraries were pooled and sequenced on an Illumina NovaSeq 6000 (University of Colorado Genomics Core) as 150bp paired-end reads.

### RNA-seq

RNA was extracted from MDBK and BL3.1 cells using the Zymo Quick RNA Miniprep Plus Kit. Monocyte and leukocyte lysates were prepared using Trizol, and RNA was extracted using the Zymo Direct-zol RNA Miniprep Kit. All RNA lysates were stored at -80 °C until processing, and single-use aliquots of extracted RNA were stored -80°C until library preparation. RNA integrity was quantified using High Sensitivity RNA TapeStation 4200. polyA enrichment and library preparation was performed using the KAPA mRNA HyperPrep Kit according to the manufacturer’s protocols. Briefly, either 200 or 500 ng of RNA was used as input. 1.5 uM single-index or unique dual-index adapters were added at a final concentration of 7 nM. Purified, adapter-ligated library was amplified for a total of 10 (500 ng) or 12 (200 ng) cycles following the manufacturer’s protocol. The final libraries were pooled and sequenced on a NovaSeq 6000 (University of Colorado Genomics Core) as 150bp paired-end reads.

### External Datasets

Publicly available data were downloaded from public repositories using fasterq-dump from the NCBI SRA Toolkit. RNA-seq datasets were obtained from PRJEB22535 and GSE116734. Whole genome sequencing datasets were obtained from PRJDB2660, PRJEB1829, PRJNA176557, PRJNA210519, PRJNA277147, PRJNA324822, PRJNA379859, and PRJNA325058. Individual sample library accession codes are available in Supplemental Table S1 and S15.

### RNA-seq analysis

Adapters and low quality reads were trimmed using BBDuk v38.05 (Bushnell sourceforge.net/projects/bbmap/) using options ‘ktrim=r k=23 mink=11 hdist=1 maq=10 qtrim=r trimq=10 tpe tbo literal=AAAAAAAAAAAAAAAAAAAAAAA’. Library quality was assessed using FastQC v0.11.8 and MultiQC v1.7 (Ewels et al. 2016), and trimmed reads were aligned to the bosTau9 (ARS-UCD1.2) assembly using HISAT2 v2.1.0 (Kim et al. 2019) with options ‘--rna-strandness RF --no-softclip’. Only uniquely aligned fragments (MAPQ >= 10) were retained using samtools v1.10. Alignments from technical replicates were merged after alignment. For visualization aligned fragments were converted to stranded, CPM normalized bigwigs using deepTools bamCoverage v3.0.1 (Ramírez et al. 2014) with options ‘--normalizeUsing CPM --ignoreForNormalization chrX chrM’. Aligned fragments were assigned to the bosTau9 refseq gene annotation in a stranded manner using featureCounts v1.6.2 (Liao et al. 2014) with options ‘-s 2 -t exon -g gene_id’, and differentially expressed genes between stimulated and unstimulated cells were called using DESeq2 v1.26.0 (Love et al. 2014). For visualization, log2 fold changes values were shrunken using the apeglm function v1.8.0 (Zhu et al. 2019).

ISGs were defined as genes with a false discovery rate of at least 0.05 and log2 fold change greater than 0. Nonresponsive genes were defined using the following cutoffs: 1) mean normalized expression >= 100, padj >= 0.90, log2 fold change >= -0.05, log2 fold change <= 0.05. Loci were collapsed to their transcriptional start site to determine relative distances from IFNG-inducible H3K27ac peaks and TEs. The venn diagram was prepared using Intervene v0.6.4 (Khan and Mathelier 2017).

### Mutant RNA-seq analysis

Mutant RNA-seq data were analyzed in a manner similar to the wildtype RNA-seq (see above) with adjustment to the differential expression analysis. Differentially expressed genes were called using DESeq2 v1.26.0 (Love et al. 2014) using a two-factor analysis incorporating both genotype (wildtype/mutant) and treatment (untreated/IFNG) information. Five clones showed editing at MER41_BT.IFNAR2, and we selected two clones (KOs #2, #3) that showed the desired deletion to define differentially expressed genes by RNA-seq. Independent mutant clones were treated as biological replicates, and reported expression differences were called comparing mutant IFNG against wildtype IFNG. As controls, we used: 1) The parent MDBK clonal line stably expressing Cas12 and 2) A MDBK/Cas12 clonal line recovered from the MER41.IFNAR2 gRNA limited dilution screen. Sanger sequencing of the deletion locus in this clone revealed evidence of minor edits at the gRNA target sites but not the intended deletion of MER41_BT.IFNAR2 (Supplemental Figure S10B). This clone was used as an effective wildtype control replicate. Two replicates were included for both controls and MER41_BT.IFNAR2 deletions. One replicate was included for all MER41_BT.LOC510185 deletions. For visualization, log2 fold changes values were shrunken using the ashr function v2.2-47 (Stephens 2017). Gene loci were collapsed to their transcriptional start site to determine relative distances to each deletion site.

### ATAC-seq analysis

Adapters and low quality reads were trimmed using BBDuk v38.05 (Bushnell sourceforge.net/projects/bbmap/) using options ‘ktrim=r k=34 mink=11 hdist=1 tpe tbo qtrim=r trimq=10’. Trimmed reads were aligned to the bosTau9 assembly (chromosomal and mitochondrial scaffolds only) using Bowtie2 v2.2.9 (Langmead and Salzberg 2012) with options ‘--end-to-end --very-sensitive -X 1000 --fr’, and only uniquely mapping reads with a minimum MAPQ of 10 were retained. Fragments aligning to the mitochondrial genome were removed, and duplicates were removed using sambamba v0.6.9. Remaining fragments were used to call ATAC-seq peaks with an FDR < 0.05 using MACS2 v2.1.1 (Liu 2014) with options ‘--keep-dup all --format BAMPE’. Scores corresponding to the fraction of reads in called peak regions (FRIP) and transcriptional start site enrichment using the complete RefSeq bosTau9 annotation were calculated to assess library quality (Supplemental Table S3). One untreated replicate was removed from the analysis as it yielded significantly fewer peaks and lower FRIP and transcriptional start site enrichment scores than other samples. For intersecting with CUT&RUN data or TEs, replicate peak files (where available) were concatenated, and peaks that fall within 100 bp of another were merged using bedtools v2.28.0. Normalized profiles corresponding to read coverage per 1 million reads were used for heatmap and metaprofile visualization, which were generated using deepTools v3.0.1 (Ramírez et al. 2014).

### CUT&RUN analysis

#### Alignment and peak calling

Adapters and low quality reads were trimmed using BBDuk v38.05 (Bushnell sourceforge.net/projects/bbmap/) using options ‘ktrim=r k=34 mink=11 hdist=1 tpe tbo qtrim=r trimq=10’. Trimmed reads were aligned to the bosTau9 assembly using BWA-MEM v0.7.15 (Li 2013), and only uniquely mapping reads with a minimum MAPQ of 10 were retained. Fragments aligning to the mitochondrial genome were removed, and alignments corresponding to replicate samples were merged. Aligned fragments from transcription factor pulldowns were subset using a 150 bp cutoff using deepTools v3.0.1 (Ramírez et al. 2014). Peak calling was performed using complete and size subsetted alignment files with MACS2 v2.1.1 (Liu 2014) in a two-step process where separate sets of peaks were called with 1) single-end options ‘--format BAM --shift=-75 --extsize=150’ and 2) paired-end option ‘--format BAMPE’. For both modes only peaks with a p-value < 0.01 were retained, and all libraries were normalized against respective IgG control libraries. Independently, peaks were called from IgG control libraries using MACS2 v2.1.1 (Liu 2014) with paired-end option ‘--format BAMPE’ for visualizing background signal enrichment. Normalized bigwigs corresponding to read coverage per 1 million reads were used for heatmap and metaprofile visualization, which were generated using deepTools v3.0.1 (Ramírez et al. 2014). Only peaks from complete (not size-subset) transcription factor libraries were used for further analyses, although size-subset bigwigs were used for genome browser screenshots.

#### Identifying IFNG-inducible CUT&RUN peaks

All merged peaks for a particular pulldown (across all replicates, untreated and IFNG-stimulated) were concatenated into a single list, and aligned fragments from each individual sample were counted for all peaks using bedtools v2.28.0. IFNG-inducible and IFNG-downregulated peaks were called using DESeq2 v1.26.0 with an FDR < 0.05 and a log2 fold change > 0 and < 0, respectively. Nonresponsive H3K27ac peaks were defined using the following cutoffs: mean normalized count >= 100 or 500, padj >= 0.90, log2 fold change >= -0.075 or -0.05, log2 fold change <= 0.075 or 0.050. For signal heatmaps, signal metaplots, and genome browser screenshots, differentially enriched H3K27ac peaks were centered on concatenated and merged (within 100 bp) ATAC peaks to improve resolution. Motif analysis was performed using XSTREME v5.4.1 (Grant and Bailey 2021) with options ‘--minw 6 --maxw 20 --streme-motifs 20 --align center’ querying against the JASPAR CORE 2018 vertebrates database (Fornes et al. 2020).

#### Enrichment near IFNG-stimulated genes

The top 750 ISGs were extracted from the larger list of 1496 ISGs sorted by descending log2 fold change. The absolute distance to the nearest ISG was determined for all 4525 IFNG-inducible H3K27ac peaks. The expected background was determined by random shuffling using bedtools v2.28.0. Additionally 4525 nonresponsive H3K27ac peaks were extracted from a larger list of 5011 peaks by random sampling. Statistical significance was determined for the first 20kb bin by Fisher’s exact test using bedtools v2.28.0. The analysis was repeated using two additional gene sets: 1) top 750 significantly downregulated genes, sorting by ascending log2 fold change and 2) randomly sampled 750 nonresponsive genes.

### Transposable element analysis

All TE analysis in this section was performed using the RepeatMasker annotations for bosTau9 available on UCSC (updated 2018-11-07). For motif analyses, binding motif position-weight matrices for STAT1 (MA0137.3 for Gamma Activated Site/GAS motif, MA0517.1 for the Interferon Stimulated Response Element/ISRE motif) were obtained from the JASPAR CORE 2018 vertebrate database (Fornes et al. 2020). Motif occurrences within TE consensus sequences or the bosTau9 cattle assembly were identified using FIMO v5.0.3 (Grant et al. 2011) using a p-value cutoff of 1⨯10^−4^ (TE heatmaps) or 1⨯10^−3^ (TE motif analysis).

#### Identifying TE-derived inducible enhancers

IFNG-inducible peaks were defined using H3K27ac, but H3K27ac-enriched regions are too broad (>1 kb) and often encompass multiple TEs. We refined peak locations by centering them on ATAC-seq summits (1 bp) contained within the H3K27ac-defined region, with the rationale that these summits represent key transcription factor binding sites within the enhancer. We then asked whether these summits overlapped an annotated TE using bedtools v2.28.0.

#### Identifying overrepresented TE families

To assess family-level enrichment, GIGGLE v0.6.3 (Layer et al. 2018) was used to create a database of all TEs in the bosTau9 genome. Merged +IFNG ATAC-seq and CUT&RUN peaks were then queried against the TE database. Results were ranked by descending Giggle enrichment score, and enriched TE families were identified according to the odds ratio, Fisher’s two-tail p-value, and number of overlaps.

#### TE heatmaps

To assess ATAC-seq and CUT&RUN normalized signal pileup, a list of 101 and 11928 TEs were subset from all 5491 MER41_BT and 362131 Bov-A2 elements, respectively. TEs were retained if they: 1) overlapped merged +IFNG ATAC-seq peaks, 2) did not fall within 5kb of another ATAC-overlapping TE, 3) did not fall within 5kb of a gene TSS (Refseq annotation downloaded from the UCSC bosTau9 assembly), and 4) overlapped a putative GAS or ISRE motif as defined by FIMO (Grant et al. 2011) with a p-value cutoff of 1⨯10^−4^. Signal from CPM normalized bigwigs was plotted over subset TEs as heatmaps using deepTools v3.0.1 (Ramírez et al. 2014).

#### TE motif analysis

For the heatmap visualization of motif presence, only MER41_BT that were 50% full-length relative to consensus were retained. BOV-A2 that were within 5% of full-length were retained. Repeat 5’ start coordinates were recalculated based on their alignment to the consensus using the RepeatMasker annotations. For all TE motif heatmaps, motif presence was plotted using putative GAS or ISRE motifs as defined by FIMO with a p-value cutoff of 1⨯10^−3^.

#### Enrichment near IFNG-stimulated genes

The top 750 ISGs were extracted from the larger list of 1496 ISGs sorted by descending log2 fold change. The absolute distance to the nearest ISG was determined for 4198 TE-derived IFNG-inducible H3K27ac peaks (centered using ATAC-seq summits as described above). The expected background was determined by randomly shuffling ATAC-centered IFNG-inducible H3K27ac peaks using bedtools v2.28.0. 3996 ATAC-centered nonresponsive H3K27ac peaks were used as an additional comparison. Statistical significance was determined for the first 20kb bin by Fisher’s exact test using bedtools v2.28.0. The analysis was repeated using two additional gene sets: 1) top 750 significantly downregulated genes, sorting by ascending log2 fold change and 2) randomly sampled 750 nonresponsive genes.

#### Re-annotation of MER41 repeats

High quality assemblies from 30 mammalian species representing 5 distinct lineages (primate, cetartiodactyla, perissodactyla, pholidota, carnivora) were annotated using RepeatMasker v4.1.0 (Smith A, Hubley R. RepeatModeler Open-1.0. In: RepeatMasker Open-4.0) with options ‘-e rmblast -s -gccalc’ using the RepeatMasker-RepBase (RMRB, release 20181026) library. We included all MER41-like subfamilies annotated in cetartiodactyla (MER41_Vpa, MER41_SS, MER41_BT, and MER41_Ttr) and included primate MER41 (MER41_Hs) as a control. For each assembly, the numbers of annotated MER41, Bov-A2, and BovB elements from 24 assemblies were visualized in a bubble plot. For MER41 and Bov-A2, only elements at least >80% (MER41) and >50% (Bov-A2) of the consensus length were considered to avoid overcounting and false positive matches. BovB elements were not filtered by length to retain copies that may have undergone 5’ truncation as a consequence of the copy-and-paste retrotransposition mechanism (Pasquesi et al. 2018). Species divergence times and phylogeny were obtained from TimeTree (Kumar et al. 2017). To determine MER41 subfamily relatedness, annotated MER41 elements across all cetartiodactyl subfamilies were collapsed into a list of 7556 elements, and only non-overlapping elements that were between 400 and 800 nt were extracted (N=990). Extracted elements were aligned using MUSCLE v3.8.1551 (Edgar 2004), and a PCA derived from the multiple sequence alignment was prepared using Jalview 2.11.1.4 (Waterhouse et al. 2009).

#### Analysis of TE syntenic conservation

Evolutionary lineage placements based on copy number were independently verified by analysis of presence or absence of syntenic MER41 subfamilies using the TE_Orthology script (https://github.com/4ureliek/TEorthology) (Kapusta et al. 2017). For each annotated cetartiodactyla assembly, all MER41 insertions that are greater than 200 bp and not nested in another annotated repeat were extracted including 100 nt 5’ and 3’ flanking sequences. Extracted MER41 elements were queried against the other 18 cetartiodactyla assemblies using BLASTN v2.7.1 (Camacho et al. 2009) with an E-value cutoff of 1⨯10^−25^. Hits are filtered based on the presence of at least 20% 5’ and 3’ flanking sequence, and the total number and number of potentially orthologous matches are recorded.

#### Detection of TE insertion polymorphisms

Publicly available whole genome, paired-end sequencing data (mean read length: 148bp) were downloaded using fasterq-dump v2.10.5. Adapters and low quality reads were trimmed using BBDuk v38.05 (Bushnell sourceforge.net/projects/bbmap/) using options ‘ktrim=r k=34 mink=11 hdist=1 tpe tbo qtrim=r trimq=10’. Trimmed reads were aligned to bosTau9 reference assembly with the bosTau5 Y chromosome (ARS-UCD1.2_Btau5.0.1Y, made available by the 1000 Bull Genomes Project) assembly using BWA-MEM v0.7.15 (Li 2013). Technical replicates were merged using Picard v2.6.0 (“Picard Toolkit.” 2019. Broad Institute, GitHub Repository. http://broadinstitute.github.io/picard/; Broad Institute), and coverage was estimated using QualiMap v2.2.1 (Okonechnikov et al. 2016).

MELT v2.1.5 (Gardner et al. 2017) was used to call TE variants. Briefly, MELT extracts discordant and split reads from alignment files, queries them against a TE consensus and fixed coordinates using a prepared TE .zip file, and outputs a VCF file that provides the genotype for each sample and variant call. TE .zip files were prepared for Bov-A2, BovB, and MER41_BT using the RepBase v24.02 consensus sequences allowing up to 10 mismatches per 100 bases. TE insertions were called following the MELT-Split pipeline using option ‘-r 150’ and supplying the estimated average coverage for each sample. TE deletions were called following the MELT-Deletion pipeline. Insertion calls were retained if they met the following criteria: within 10% of full length (Bov-A2 only), ASSESS > 3, SR > 3, no null genotypes in any sample. Deletion calls were retained as long as they were within 10% of full length (Bov-A2 only) and exhibited no null genotypes in any sample. Filtered and insertion variant calls were aggregated. For both the PCA and histogram, aggregated variant call genotypes were collapsed by whether or not they had a single allele supporting the variant. The PCA of the aggregated TE variant genotype calls was performed using PCAtools v2.4.0 (Blighe K, Lun A (2021). *PCAtools: PCAtools: Everything Principal Components Analysis*. R package version 2.4.0, https://github.com/kevinblighe/PCAtools.). Filtered deletion calls were called as being epigenetically marked if they overlapped any raw ATAC-seq or H3K27ac, POL2RA, STAT1, or pSTAT1 peak, and the absolute distance from each to all gene transcriptional start sites within 250kb were determined using bedtools (Supplemental Table S13). Alignments over filtered deletions were visualized using Samplot v1.1.6 (Belyeu et al. 2021) with option ‘-d 100’.

### LOC510185 sequence analysis

The inferred protein sequence for LOC510185 (RefSeq accession XP_010803655.1) and protein sequences for bovine (UniProt accession P14784) and human (F1N409) IL2RB were aligned using MUSCLE v3.8.1551 (Edgar 2004). The resulting alignment was annotated according to homology to human IL2RB according to UniProt.

## DATA ACCESS

All raw and processed sequencing data generated in this study have been submitted to the NCBI Gene Expression Omnibus (GEO) with accession number GSE185082.

## Code availability

UCSC Genome browser sessions and all code will be available at https://github.com/coke6162/bovine_TE_evolution by the time of publication.

## ACKNOWLEDGEMENTS

We thank M. Heaton for assistance obtaining genomic sequences, S. Bierman for technical assistance obtaining primary bovine cells, and USMARC cattle operations for animal handling. We thank the University of Colorado Genomics Shared Resource and BioFrontiers Computing core for technical support during this study. This study was supported by the NIH (1R35GM128822), the Alfred P. Sloan Foundation, the David and Lucile Packard Foundation, and the Boettcher foundation.

## Author contributions

C.J.K. and E.B.C conceived and designed the study and wrote the paper. C.J.K. performed all experiments and computational analyses. C.C-M contributed reagents and samples, and supervised the study.

## SUPPLEMENTARY MATERIAL

**Supplementary Fig. S1**

Differentially expressed genes from BL3.1, monocyte, leukocyte, alveolar macrophage, and bone marrow derived macrophages.

**Supplementary Fig. S2**

Mean normalized counts for *CIITA* and *TLR4* from wild type RNA-seq.

**Supplementary Fig. S3**

Genome browser view of *TLR4* locus.

**Supplementary Fig. S4**

Signal enrichment over IFNG-downregulated and nonresponsive H3K27ac peaks.

**Supplementary Fig. S5**

Absolute distances from IFNG-inducible H3K27ac to the nearest IRG and nonresponsive gene.

**Supplementary Fig. S6**

Absolute distances from TE-derived IFNG-inducible H3K27ac to the nearest ISG, IRG, and nonresponsive gene.

**Supplementary Fig. S7**

Family-level TE enrichment for IFNG-downregulated H3K27ac, nonresponsive H3K27ac, IFNG ATAC, IFNG-inducible POL2RA, and IFNG-inducible pSTAT1 peaks.

**Supplementary Fig. S8**

Inferred homology between IL2RB and LOC510185.

**Supplementary Fig. S9**

Validation of MER41_BT.IL2RB deletions

**Supplementary Fig. S10**

Validation of MER41_BT.IFNAR2 deletions

**Supplementary Fig. S11**

Mean normalized counts for *IFNGR2* from MER41_BT.IFNAR2 RNA-seq.

**Supplementary Fig. S12**

Characterization of predicted Bov-A2 and BovB variants as a function of genotype and variant plots for Bov-A2.BICC1 and Bov-A2.SNX19.

**Supplementary Table S1 (separate file)**

Wild type RNA-seq alignment statistics and DESeq2 results for MDBK cells, BL3.1 cells, monocytes, leukocytes, alveolar macrophages, and bone marrow derived macrophages.

**Supplementary Table S2 (separate file)**

gProfiler gene ontology results for wild type RNA-seq upregulated genes.

**Supplementary Table S3 (separate file)**

MDBK ATAC-seq alignment statistics.

**Supplementary Table S4 (separate file)**

MDBK CUT&RUN alignment statistics and DESeq2 results.

**Supplementary Table S5 (separate file)**

CUT&RUN motif enrichment using XSTREME.

**Supplementary Table S6 (separate file)**

TE-derived IFNG-inducible, IFNG-downregulated, and nonresponsive H3K27ac peaks.

**Supplementary Table S7 (separate file)**

Family-level TE enrichment using GIGGLE.

**Supplementary Table S8 (separate file)**

Absolute distances for IFNG-inducible MER41_BT to the nearest MDBK ISG.

**Supplementary Table S9 (separate file)**

Alignment statistics and DESeq2 results from MER41_BT.IL2RB and MER41_BT.IFNAR2 deletions in MDBK cells.

**Supplementary Table S10 (separate file)**

Summary of assemblies used in RepeatMasker analysis.

**Supplementary Table S11 (separate file)**

MER41 synteny analysis using TEOrthology.

**Supplementary Table S12 (separate file)**

Filtered Bov-A2, BovB, and MER41_BT insertions and deletions as called using MELT.

**Supplementary Table S13 (separate file)**

Absolute distances for epigenetic-marked Bov-A2 to nearest gene transcriptional start site (TSS)

**Supplementary Table S14 (separate file)**

Reagents and sequences used in this study.

**Supplementary Table S15 (separate file)**

Identifying information for all publicly available whole genome sequencing data analyzed in this study.

